# The folded X-pattern is not necessarily a statistical signature of decision confidence

**DOI:** 10.1101/426635

**Authors:** Manuel Rausch, Michael Zehetleitner

## Abstract

Recent studies have traced the neural correlates of confidence in perceptual choices using statistical signatures of confidence. The most widely used statistical signature is the folded X-pattern, which was derived from a standard model of confidence assuming an objective definition of confidence as the posterior probability of making the correct choice given the evidence. The folded X-pattern entails that confidence as the subjective probability of being correct equals the probability 0.75 if the stimulus in neutral about the choice options, increases with discriminability of the stimulus in correct trials, and decreases with discriminability in incorrect trials. Here, we show that the standard model of confidence is a special case in which there is no reliable trial-by-trial evidence about discriminability itself. According to a more general model, if there is enough evidence about discriminability, objective confidence is characterised by different pattern: For both correct and incorrect choices, confidence increases with discriminability. In addition, we demonstrate the consequence if discriminability is varied in discrete steps within the standard model: confidence in choices about neutral stimuli is no longer .75. Overall, identifying neural correlates of confidence by presupposing the folded X-pattern as a statistical signature of confidence is not legitimate.

## Introduction

Confidence is a metacognitive evaluation of decision making: Each choice can be accompanied by some degree of confidence that the choice is correct. In neuroscience, confidence has become a flourishing research topic, uncovering the underlying neural mechanisms in humans [1–6] as well as non-human animals [7–13]. A major obstacle to the scientific study of confidence is the inherently subjective nature of the psychological construct of decision confidence. Therefore, a large amount of recent research on confidence has been inspired by a novel approach that formalizes confidence mathematically as an objective statistical quantity [14, 15]. This formalization defines confidence as the belief that a choice is correct [16]. From a Bayesian perspective, beliefs are best formalised as probabilities [17, 18]. Decision confidence in this formalization is the posterior probability of being correct given the evidence [16, 19]. Several predictions about objective confidence have been formally derived from the model to which we subsequently refer to as the standard model of confidence [7,14,15]: First, the average objective confidence in correct choices increases as a function of the discriminability of the stimulus. Second, the average confidence in incorrect choices decreases with discriminability. Finally, when the stimulus is neutral about the choice options, confidence is exactly 0.75. The overall pattern, which we refer to here as folded X-pattern [20], has been dubbed a “statistical signature of confidence” [14, 21]. Given that the folded X-pattern follows objectively from the posterior probability of being correct, it has been argued that when the folded X-pattern is detected in another behavioural, neural, or physiological variable, that variable should be considered a correlate of confidence [7,14,15,22]. Thus, it is frequently used to empirically identify correlates of decision confidence [7,8,10,22–25]. Nevertheless, a recent study suggested that the Bayesian calculation of the posterior probability of being correct does not necessarily imply the folded X-pattern [26]. The inverse is also not correct as the folded X-pattern does not necessary imply the Bayesian calculation of confidence [27–29].

Here, we show that the folded X-pattern is no longer expected when confidence is informed by a trial-by-trial representation of discriminability. When objective confidence is calculated from a model of confidence which is more general in the sense that it includes a representation of discriminability, the folded X-pattern occurs only as a special case when the evidence about the discriminability of a specific stimulus is not reliable. When there is accurate information about the discriminability of a stimulus, confidence tends to increase as a function of discriminability in correct and incorrect trials, which is why we refer to this pattern as the double increase pattern.

### The standard model of confidence

The standard model of confidence is depicted in Fig 1. According to Sanders et al. [14], when an observer is presented with a stimulus and asked to make a choice ϑ ∈ {−1,1} about the stimulus, the stimulus d is a continuous variable that differentiates between the two options of ϑ. Negative values of d mean that observers should choose ϑ = −1; d = 0 means no objective feature of the stimulus suggests any of the two options, and positive values indicate that observers ought to choose ϑ = 1. As the sign of d determines what response observers ought to give, we refer to the sign of the stimulus as identity I. The absolute value of |d| is referred to as discriminability: The greater is the distance between d and 0, the easier is the choice. The accuracy of the choice A is 1 if I and ϑ are the same, and 0 otherwise. However, observers cannot perceive d directly, instead, the choice is based on noisy sensory evidence eI (referred to as percept by Sanders et al.), which can be considered an estimate of 1. d. The most frequent approach is to model eI as a random sample from a Gaussian with a mean of d, while ϑ is modelled as a deterministic function of d. Finally, given that observers know the distributions from which d and eI are sampled, the posterior probability of a correct choice given the sensory evidence eI and the choice ϑ can be calculated using Bayes’ theorem (see S1 Appendix).

**Fig 1.**
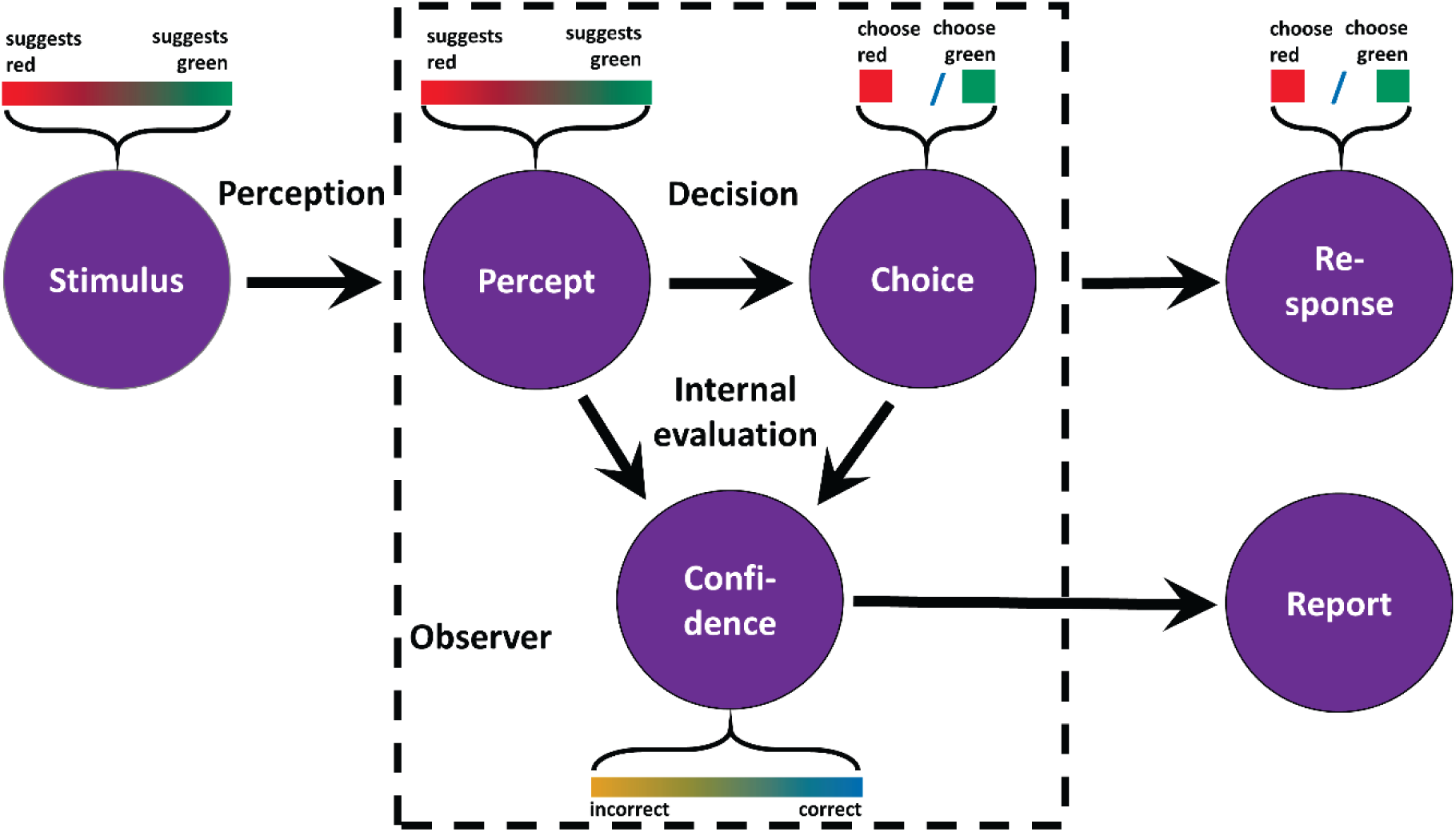
The standard model of confidence. The stimulus objectively supports the choice options “red” and “green” to varying degrees. As perception is noisy, the percept is a corrupted representation of the degree to which the stimulus favours a specific choice option. Confidence is the probability of making the correct choice given percept and choice.

The standard model has been presupposed to derive the folded X-pattern [14, 15], although different aspects of the folded X-pattern come with specific additional assumptions: First, confidence in choices about neutral evidence is .75 only if the distribution of the stimulus is uniform and yields choice accuracies spanning from 0.5 to 1, and if sensory evidence is sampled from a symmetric distribution with a single peak centred on the stimulus, and if choice is deterministic [14,15,26]. Second, the decrease of confidence in incorrect choices presupposes that the observer is not provided with any information about the discriminability of the stimulus at the level of single choices [15]. Although the Bayesian calculation of the probability of being correct implies knowledge of the distribution from which d is sampled, knowledge the distribution of d only implies that observers know the probability of the degrees of discriminability across the experiment. For each specific choice however, the standard model assumes that observers do not possess any knowledge what the discriminability of the stimulus is over and above the distribution from which d is sampled.

### The general model of confidence

The general model of confidence extends the standard model by including the possibility that observers perceive or infer the discriminability of the stimulus on the level of single choices. For example, when a driver in heavy rain needs to discern if a traffic light is green or red, the driver might not only be unsure because their colour percept is ambiguous, but they might also be cautious because they see or know their view is hindered by rain. Analogous to traffic lights and rain, many psychophysical experiments do not manipulate the stimulus as one independent variable; instead, two features of the stimulus are varied across the experiment. Therefore, the general model of confidence (see Fig 2) considers identity I and discriminability d as two independent aspects of each single stimulus: The identity, which in each trial can be either −1 or 1, is the variable in the external world that determines which of the choice options is correct. The model generates a choice ϑ about the identity I of the stimulus. For example, the stimulus could be red or green, and participants need to make a choice accordingly. Choices are correct when I and ϑ are both either −1 or 1.

**Fig 2.**
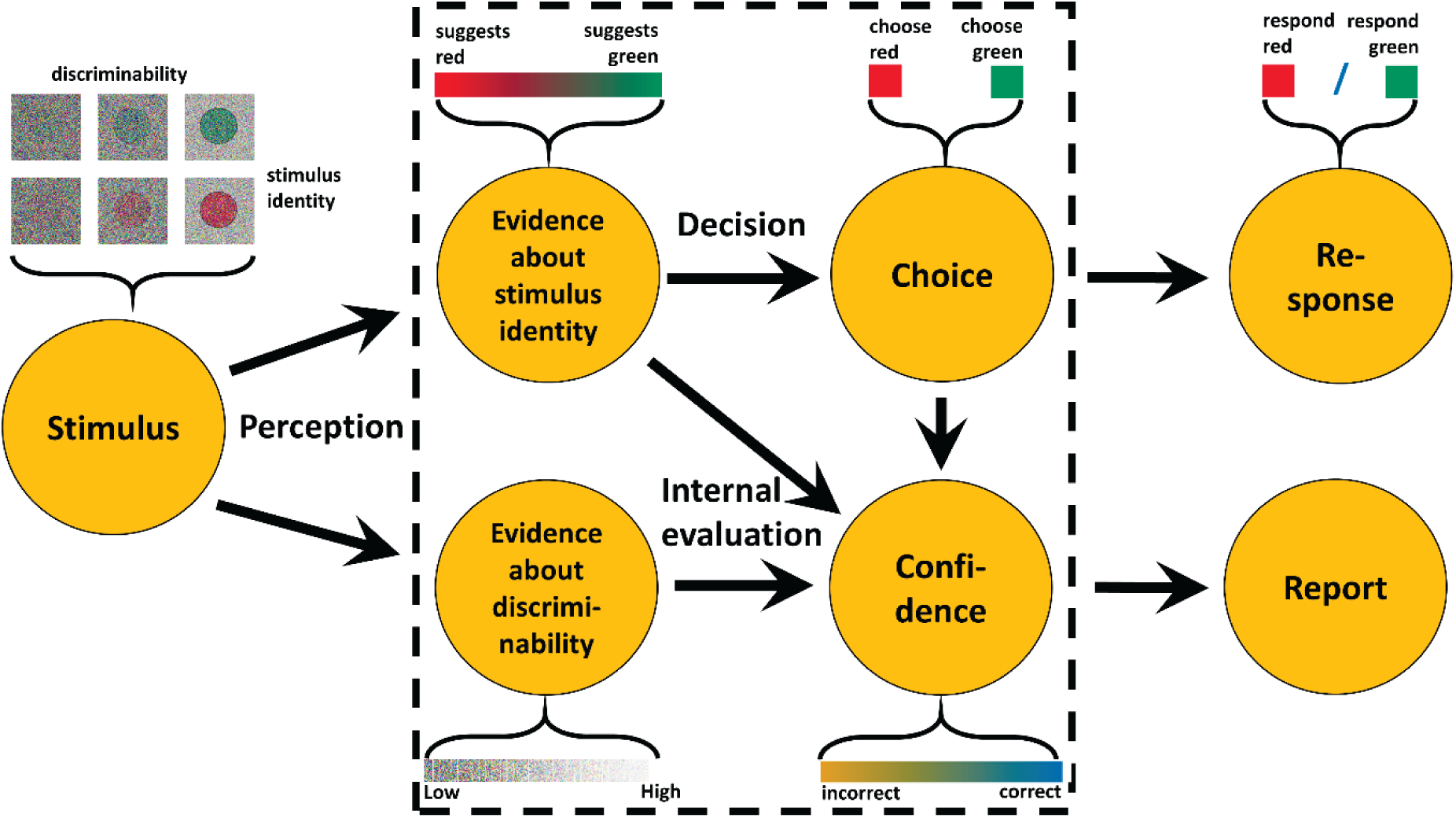
The general model of confidence. The general model is a generalization of the standard model. In many psychophysical experiments, the stimulus varies in two aspects: stimulus identity (symbolized here as red and green colour patches) and discriminability (symbolized here by the noise dots). In the general model, the stimulus generates two internal variables: the evidence about the stimulus identity, a continuous variable that differentiates between the possible identities, and evidence about the discriminability. Objective confidence about the correctness of the choice is based on evidence about the identity as well as evidence about discriminability.

Discriminability d is the variable in the external world that determines how easy/difficult the choice is. For instance, many experiments manipulate contrast, presentation time, or luminance orthogonally to stimulus identity I. According to the general model, observers in each single trial obtain sensory evidence about *both* aspects of the stimulus, i.e. there is sensory evidence for identity e_I_, and evidence for discriminability e_d_. While e_I_ depends on I and on d, e_d_ depends only on d, but not on I. To represent that observers’ do not have direct access to I and d, e_d_ is sampled from a Gaussian distribution whose mean depends on d, and e_I_ is sampled from a Gaussian whose mean depends on I and on d. The posterior probability of a correct choice given and the choice ϑ can again be calculated based on Bayes’ theorem (see S2 Appendix).

There are at least two possibilities why in an experimental situation, evidence about discriminability e_d_ may exist separately from the evidence about the identity e_I_: First, when stimuli with different degrees of discriminability are not presented in random sequence, for example when discriminability is constant within one block of the experiment, observers can infer the discriminability of the present stimulus. A second possibility is that observers in many cases are able to perceive discriminability directly: Within the visual system, there is not only sensory evidence about the choice-relevant stimulus feature I, but also sensory evidence about other features of the stimulus, irrelevant to the current choice [30, 31]. For example, in a masked orientation task, observers may estimate the discriminability not only by their percept of the orientation, but also by their percept of the shape, texture, or presentation time of the stimulus, even when these features are not explicitly manipulated by the experimenter [32]. All sensory evidence irrelevant to the current choice can be used as evidence about the discriminability as long as it is correlated with discriminability.

Why is confidence not exclusively based on sensory evidence dependent on the choice-relevant features of the stimulus if decision confidence is calculated objectively, but also on evidence for the quality and reliability of perception itself? The key fact is that confidence as the posterior probability that the choice is correct given the evidence is only objective if it includes all information that is dependent on the stimulus. Given confidence is objective only if all evidence available is used, and if e_d_ exists in a specific task, it follows that objective confidence should be based on e_d_, too.

### Rationale of the present study

In the present study, we used Monte Carlo simulations to trace the statistical patterns of optimal confidence calculated as the posterior probability of being correct given the evidence. Our simulations were based on the standard model as well as on the general model, which extends the standard model by assuming that observers on single trial basis obtain evidence about the discriminability of the stimulus. Based on the general model, we also examined the impact of the reliability of evidence about discriminability on the statistical pattern of confidence. Finally, we examined if relying confidence on evidence about discriminability is a beneficial strategy, or if it is an example of a suboptimal mental shortcut to the probability of being correct [6,27,33–36], i.e. a heuristic [37, 38].

## Results

### Standard model

Fig 3 shows the patterns of confidence obtained from simulations based on the standard model. Only two of the three postulated features of the folded X-pattern consistently follow from the standard model: Independent of the distribution of discriminability |d|, confidence in correct choices always increases as a function of discriminability, and confidence in incorrect choices always decreases with discriminability. However, when the stimulus is neutral about the choice options, confidence is .75 only when |d| is sampled from a continuous uniform distribution that includes high discriminability (see Fig 3f). When |d| is sampled from a discrete uniform distribution (Fig 3a-c) or a gamma distribution (Fig 3d, e), or when the continuous uniform distribution does not support high discriminability (Fig 3a, b), confidence in choices about neutral stimuli is not .75.

**Fig 3.**
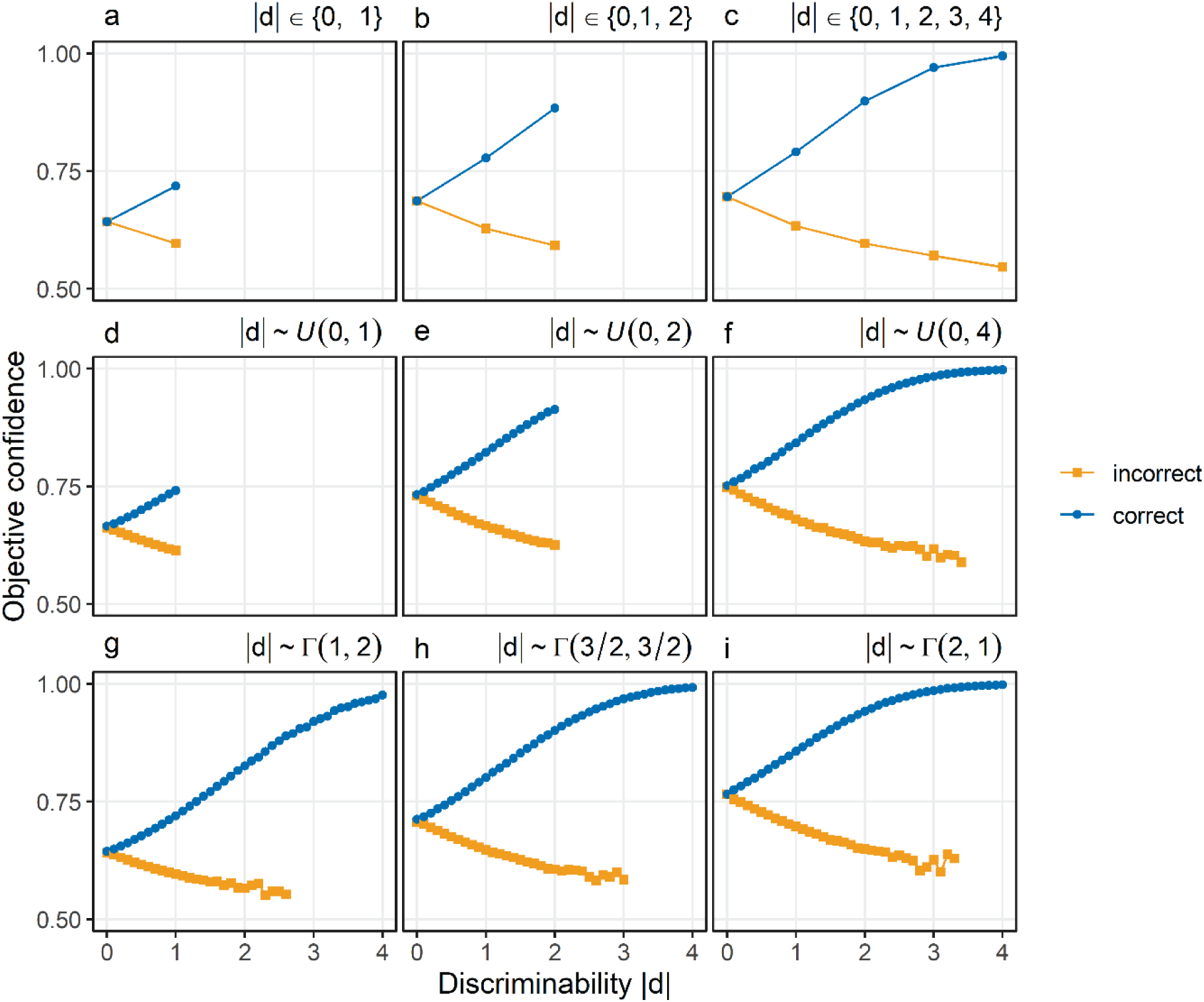
Objective confidence given the standard model of confidence. Confidence (y-axis) is shown as a function of discriminability (x-axis) in correct choices (blue) and incorrect choices (orange). Different panels show different distributions from which discriminability was sampled. Panels a-c: Discrete uniform distributions. Panels d-f: Continuous uniform distributions. Panels g-i: Gamma distributions. In all simulations, the percept eI was sampled from a normal distribution with a mean equal to the stimulus d and a standard deviation σ_I_ of 1.

Fig. 4 shows the effect of the number of possible values for |d| with a constant maximal level for |d|, assuming a finite number of possible values as well as an equal probability of each value. When there are only few possible values for |d|, confidence in choices with neutral evidence is below .75 (see Fig 4a, b). Only when the number of discrete possible values increases – and thus the distribution which |d| is sampled from becomes more similar to a continuous uniform distribution - confidence in choices about neutral evidence becomes close to .75 (see Fig 4c, d).

**Fig 4.**
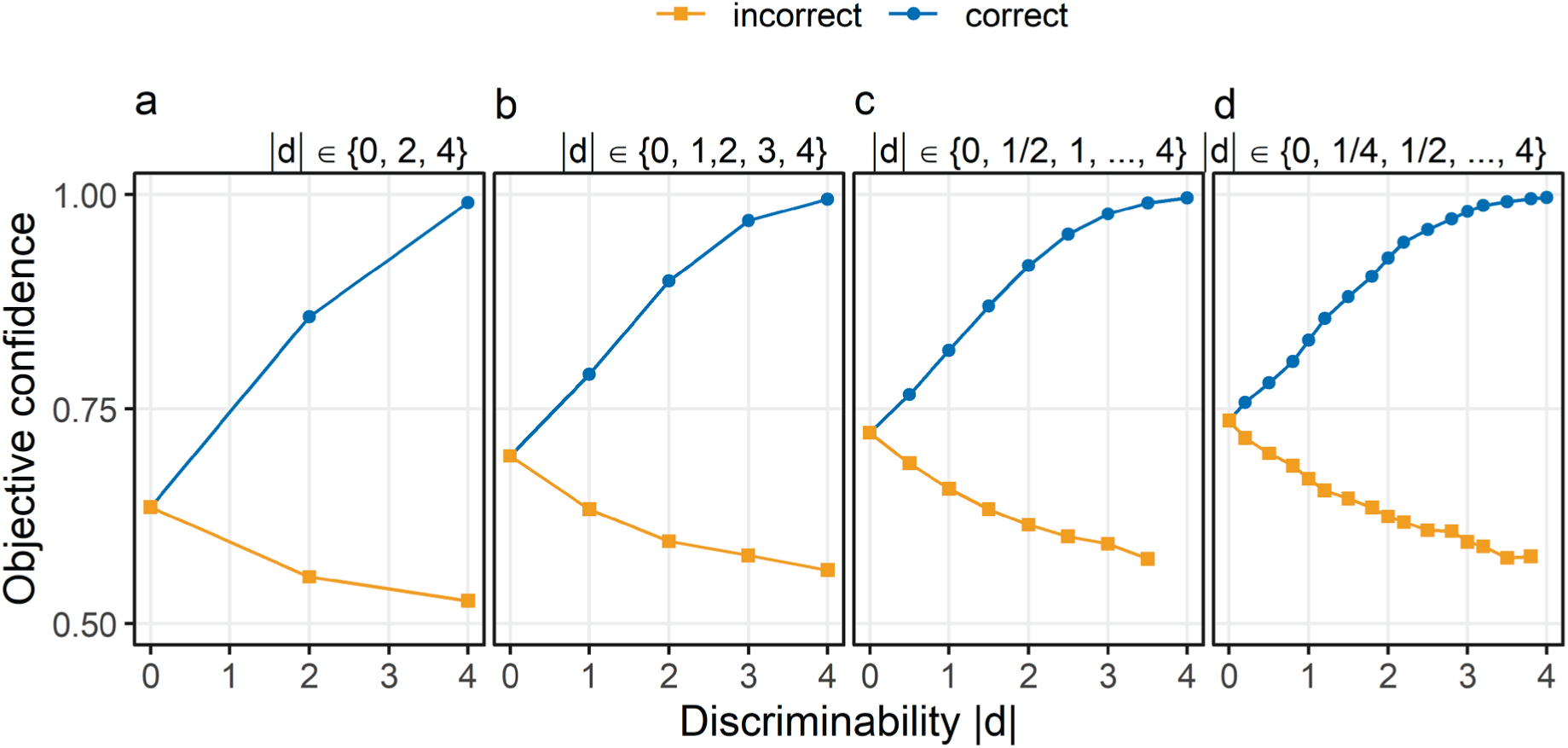
Objective confidence in standard model depending on the number of different levels of discriminability. Confidence (y-axis) is shown as a function of discriminability (x-axis) in correct choices (blue) and incorrect choices (orange). Different panels show different numbers of levels of discriminability |d|, sampled from discrete uniform distributions. Possible values of |d| are 0, 2, and 4 (Panel a), 0, 1, 2, 3, and 4 (Panel b), 0, ½, 1, …, 4 (Panel d or 0, ¼, ½, …, 4 (Panel d). The percept e_I_ was sampled from a normal distribution with a mean equal to the stimulus d and a standard deviation σ_I_ of 1.

Fig 5 shows the patterns of confidence assuming only two equally probable values of |d|. In this case, confidence in choices about neutral evidence is not .75, irrespective of whether neutral stimuli are paired with hard decisions (see Fig 5a, b), or with easier decisions (see Fig 5c, d).

**Fig 5.**
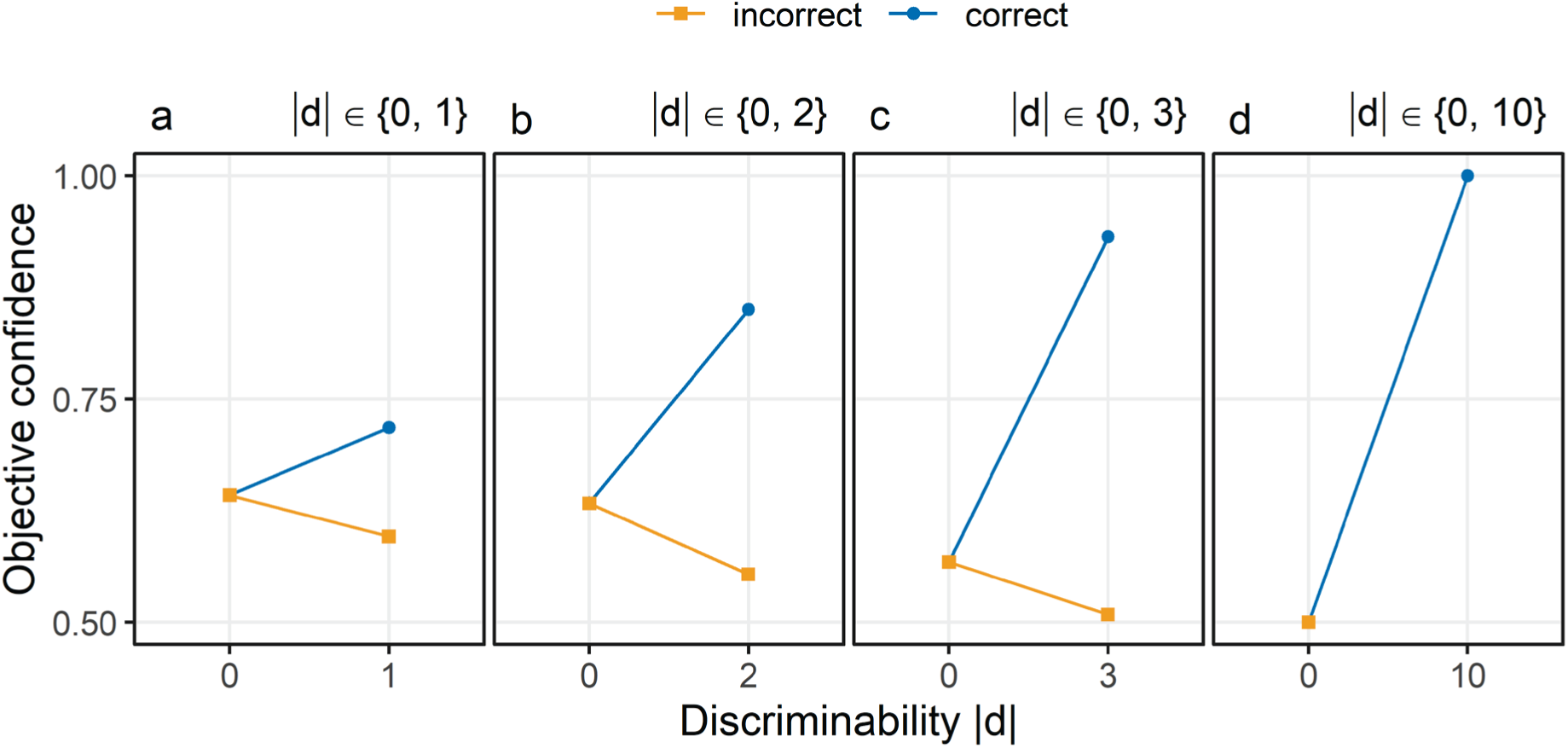
Objective confidence in the standard model if discriminability is either 0 or maximal. Discriminability |d| was always sampled from discrete uniform distributions with only two values. One of the two possible values was always 0, indicating neutral stimuli with respect to the choice options. The second possible value of |d| was 1 (Panel a), 2 (Panel b), 3 (Panel c), or 10 (Panel d). Confidence (y-axis) is shown as a function of discriminability (x-axis) in correct choices (blue) and incorrect choices (orange). The percept eI was sampled from a normal distribution with a mean equal to the stimulus d and a standard deviation σ_I_ of 1.

### General model

What is the pattern of confidence expected from the general model? As can be seen from Fig 6, the general model is compatible with both the folded X-pattern and the double increase pattern. When σ_d_ is small and thus the evidence about discriminability is reliable (see Fig 6, a1-a9), confidence approaches 0.5 when discriminability is 0. In addition, confidence increases with discriminability for both in correct choices well as in incorrect choices, i.e. confidence is characterised by what we refer to as the double increase pattern. These patterns are the same across different distributions of discriminability (Fig 6, different rows). When σ_d_ is large and thus there is only corrupted evidence about discriminability (see Fig 6, d1-d9), the pattern of confidence is the same as for the standard model (cf. Fig 3). When σ_d_ increases (see Fig 6, b1-b9, c1-c9), confidence in choices about stimuli with d = 0 increases. Additionally, when σ_d_ increases, the correlation between discriminability and confidence in incorrect choices becomes more negative, eventually switching sign from positive to negative.

**Fig 6.**
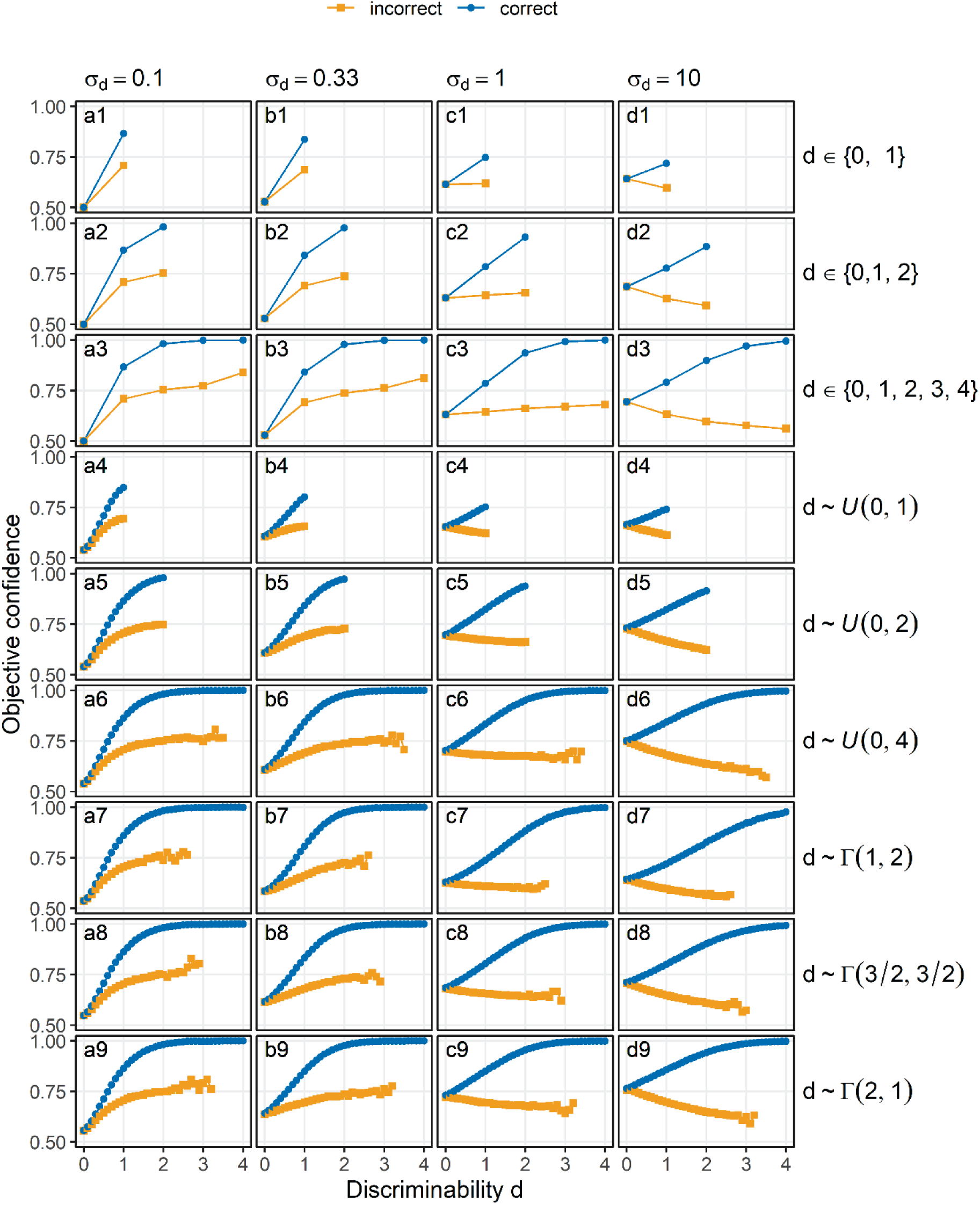
Objective confidence according to the general model of confidence. The sensory evidence about the identity e_I_ and evidence about discriminability e_d_ were both sampled from normal distributions, with standard d_ev_iations σ_I_ = 1 and σ_d_ varying across columns. Confidence (y-axis) is shown as a function of discriminability (x-axis) in correct trials (blue) and incorrect trials (orange). Panels a1-a9: σ_d_ = 0.1. Panels b1-b9: σ_d_ = 0.33. Panels c1-c9: σ_d_ = 1. Panels d1-d9: σ_d_ = 10. Different rows indicate different distributions of discriminability within the simulated experiments. Rows 1-3: Discrete uniform distributions, rows 4-6: continuous uniform distributions, rows 7-9: Gamma distributions.

### Accuracy of confidence

Accuracy of confidence was assessed by the information entropy of choice accuracy conditioned on confidence H(A|c). The information entropy is a measure of prediction error motivated by the free energy principle [39]: H(A|c) reflects the uncertainty with respect to choice accuracy given confidence; if choice accuracy is perfectly specified by confidence, H(A|c) will be zero. Fig 7 compares H(A|c) between confidence based on evidence about the identity e_I_ only and confidence based on evidence about the identity e_I_ and evidence about discriminability e_d_. The assumption that confidence is based exclusively on evidence about e_I_ is equivalent to the standard model. Fig 7 shows that when the standard deviation of the evidence about discriminability σ_d_ is low, confidence based on e_I_ and e_d_ is associated with a lower information entropy of accuracy conditioned on confidence than confidence based on e_I_ alone. This means that when there is an accurate estimate of discriminability, confidence that takes the evidence about discriminability into account is associated with a smaller prediction error than confidence ignoring evidence about discriminability. For larger values of σ_d_, H(A|c) is the same between confidence based on e_I_ and e_d_ and confidence based on e_I_, meaning that there is no longer a benefit of the estimate of discriminability when the estimate was too noisy. Importantly, even when σ_d_ is very large, there is never a case when confidence based on e_I_ and e_d_ is worse than confidence based solely on e_I_.

**Fig 7.**
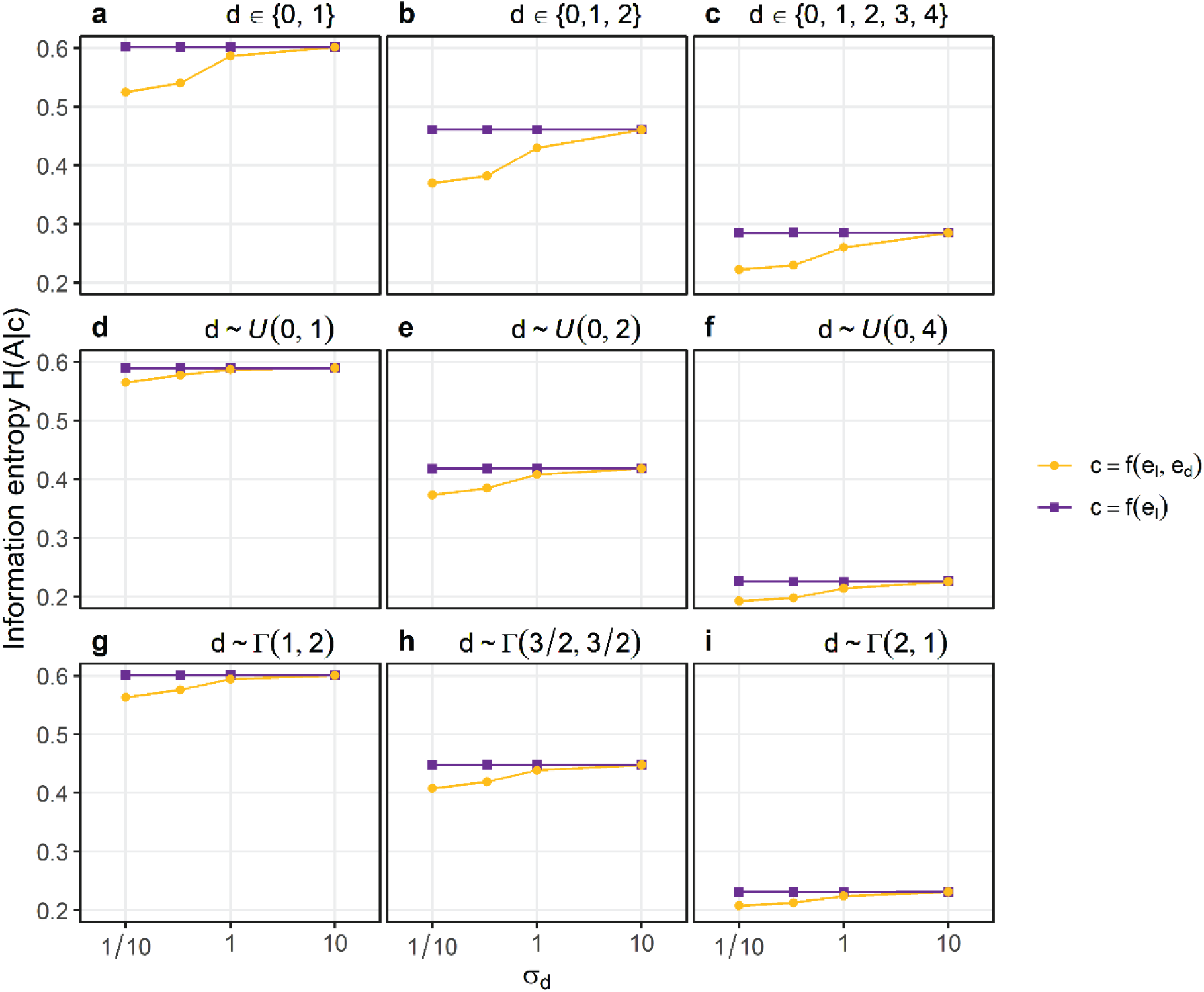
The information entropy of choice accuracy conditioned on confidence H(A|c). The noise parameters of the estimate of discriminability σ_d_ is displayed on the x-axis. Different panels indicate different distributions of discriminability within the simulated experiments. Panels a-c: Discrete uniform distributions. Panels d-f: Continuous uniform distributions. Panels g-i: Gamma distributions. Violet symbols indicate H(A|c) when confidence is calculated exclusively based on sensory evidence about the identity of the stimulus e_I_. Orange symbols indicate H(A|c) when confidence is calculated based on evidence about the identity of the stimulus e_I_ and on evidence about discriminability e_d_. The standard deviation of evidence about the identity σ_I_ was set to 1.

## Discussion

The present study showed that the objective calculation of confidence does often not imply the folded X-pattern. When there is sufficient evidence about discriminability as predicted by the general model, the correlation between discriminability and confidence in incorrect trials is positive, not negative. Even if there no evidence about discriminability, confidence in choices about neutral stimuli is not .75 unless discriminability is sampled from a continuous uniform distribution with high maximal discriminability. We also showed by simulations that if observers make optimal use of the evidence, and if evidence about discriminability is available, then confidence depends on evidence about discriminability.

The observation that the Bayesian calculation of confidence does not always imply the folded X-pattern corroborates the results of a previous study [26]. Adler and Ma showed that the folded X-pattern depends on the distribution from which the stimulus is sampled. Specifically, confidence in incorrect choices no longer decreases with discriminability if stimuli are only probabilistically related to which choice observers ought to make. Likewise, confidence in neutral events is .75 only if the width of the stimulus distribution is quite large compared to the noise in perception. The present study shows that there are at least two more cases where confidence is not expected to follow the folded X-pattern. First, when discriminability does not vary continuously but in a small number of discrete steps, optimal confidence in choices about neutral events is not .75. Notably, previous studies assuming the folded X-pattern typically relied on discrete manipulations of discriminability. Second, when observers can perceive or infer discriminability on a single trial level with sufficient accuracy, objective confidence follows the double increase pattern.

In summary, these observations imply that blind reliance on the folded X-pattern potentially leads to false conclusions. Identifying correlates of confidence by a priori presupposing the folded X-pattern is not advisable because objective confidence may not show the expected properties. Likewise, it is also not advisable to infer the computational principles underlying observed confidence judgments based on statistical signatures alone, because various different models are able to recreate the folded X-pattern [26,28,29,32], just as the double increase pattern [26,32,40]. Importantly, both the folded X-pattern and the double increase pattern are compatible with Bayesian computation of confidence, which is why model fitting is necessary to ascertain which model is the generative model of the data [26].

### Why should sensory evidence parallel to the choice improve objective confidence?

The double increase pattern has been regarded as indicative of a suboptimal mental shortcut to the probability of being correct [33], i.e. a heuristic [37, 38]. However, as evidence about discriminability in fact decreases the prediction error of confidence, the double increase pattern may in some cases indicate optimal, not suboptimal calculation of confidence.

Too see why it is necessary to include e_d_ in the calculation of objective confidence, we can look at the formula of posterior probability of the identity according to the general model (see S2 Appendix for the derivation):

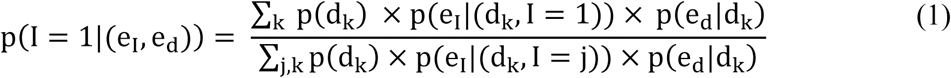

In formula (1), I represents the identity of the stimulus, e_I_ the evidence about the identity, d discriminability, and e_d_ the is the evidence about discriminability. As can be seen from the formula, evidence about the discriminability e_d_ is needed to calculate the objective posterior probability given the evidence. This means if observers make optimal use of the evidence, and if evidence about the discriminability e_d_ is available, e_d_ ought to be included into the calculation of the posterior probability of the identity and hence confidence.

Now, to get some intuition why it is optimal to include e_d_ in the calculation of confidence, let us look at formula (1) more closely. The Bayesian computation of the posterior probability divides the likelihood of the evidence about the identity e_I_ given the identity 1 (the terms in the numerator) by the sum of the likelihood of e_I_ given I = −1 and the likelihood of e_I_ given I = 1 (the terms in the denominator). Calculating the likelihood of e_I_ requires knowledge of the distribution from which e_I_ is sampled. However, according to the model, e_I_ is sampled from a Gaussian whose mean not only depends on I, but also on d. For this reason, the likelihood of e_I_ given I is calculated by multiplying the prior probability of a specific level discriminability p(d_k_) with the likelihood of e_I_ given the level discriminability and the identity p(e_I_|d_k_, I), and summing these terms across all levels of discriminability. Conceptually, these terms imply a consideration how plausible *e*_*I*_ is given the identity and given the level of discriminability, weighted by the plausibility of that level of discriminability. These terms are summed over all possible values of discriminability. The product of p(d_k_) and p(e_I_|d_k_, I) represents the case of the standard model: Observers know how plausible each degree of discriminability is across the experiment, and based on that prior information, they evaluate the plausibility of e_I_. The novel feature of the general model is the inclusion of the probability of evidence about discriminability given discriminability p(e_d_|d_k_). Conceptually, p(e_d_|d_k_) implies the evaluation how plausible the level of discriminability is based on the evidence about the discriminability. As can be seen in the formula, p(e_d_|d_k_) is multiplied with p(d_k_) and p(e_I_|d_k_, I). Thus, in the general model, observers attach a weight to p(e_I_|d_k_, I) not only based the prior knowledge of the distribution of discriminability within the experiment, but they also evaluate the plausibility of each degree of discriminability based on sensory evidence about the discriminability. Thus, evidence about the discriminability improves the efficiency of the evaluation of e_I_ because evaluating the plausibility of p(e_I_|I) requires knowledge about d, and some additional information about the discriminability is better than the prior distribution alone. If p(e_d_|d_k_) is the same across all levels of discriminability, the general model makes the same predictions as the standard model; conceptually, identical p(e_d_|d_k_) across all levels of discriminability represents the case when there is no information about discriminability on a single trial basis.

### Empirical support for folded X- and the double increase pattern

What is the empirical evidence concerning the two statistical patterns of confidence? Several previous experiments were indeed in accordance with the folded X-pattern. In an auditory discrimination task [14], a general knowledge task [14], as well as a visual two-alternative forced choice tasks [41], confidence increased with discriminability in correct trials, decreased with discriminability in incorrect trials, and was medium when stimuli could not be distinguished. The folded X-pattern was also consistent with rats’ willingness to wait for reward in an odour discrimination task [7, 24], which can be seen as a marker of confidence in non-humans.

However, six other studies based on human observers were not consistent with the folded X-pattern, and three of these studies revealed the double increase pattern instead. In two random dot motion discrimination tasks, coherence of motion was positively, not negatively, associated with confidence in incorrect trials [42, 43]. Likewise, in a masked orientation discrimination task, confidence in incorrect trials increased with stimulus-onset-asynchrony as well [32]. Two studies revealed a relationship between confidence in incorrect trials and discriminability that was essentially flat. In a second masked orientation discrimination task, in which observers’ confidence was assessed by asking observers on which of two subsequent orientation judgments they were willing to bet, confidence in incorrect trials was approximately constant across levels of stimulus contrast [44]. Moreover, in a low-contrast orientation discrimination task, the average confidence in incorrect trials was approximately constant across task difficulty levels [45]. Finally, in a discrimination task about the average orientation of a sequence of oriented Gabor patches, one subset of observers showed the folded X-pattern and another subset the double increase pattern [33], although the interpretation of the inverse variability of sequence of oriented Gabor patches as discriminability is controversial [26].

Overall, these studies suggested that the folded X-pattern is by no means universal. Although there is empirical support for the folded X-pattern in some experiments, in other experiments the pattern is just opposite to what has been considered as the signature of confidence.

How can the differences between those studies be explained? One possibility is that some experimental tasks allow observers to estimate the discriminability on a single trial basis, as predicted by the general model: Strikingly, all studies that reported an increase of confidence and incorrect choices with discriminability were based on psychophysical tasks where the stimulus was composed out of one feature that defined the response as well as an orthogonal manipulation of discriminability: In the random dot motion discrimination tasks, participants responded to the direction of motion, and the discriminability was manipulated by the coherence of the motion signal [42, 43]. Likewise, in the masked orientation task, the identity of the stimulus was defined by the orientation of the stimulus, while discriminability was manipulated by the time between stimulus onset and mask onset [32]. In contrast, those studies that observed that confidence in incorrect choices decreased with discriminability all aimed to vary the evidence more directly by using stimulus material providing different mixtures of evidence to the observer: The auditory discrimination experiment delivered click streams to both ears of the observers, and participants had to indicate which click rate was faster. Importantly, evidence was varied by the ratio between click frequencies in the two streams [14]. Likewise, the general knowledge task required observers to decide which of two countries had a greater population, with discriminability defined as the log ratio of the population size of the two countries [14]. Finally, participants in one of the two visual two-alternative forced choice tasks indicated which of two presented textured stimuli showed had un unequal amount of white and black squares. The difficulty of the task was varied by the proportion of white to black squares [41]. In all these tasks, the stimulus consisting of mixtures of evidence about the identity might make it more difficult to estimate discriminability.

An alternative explanation for the differences between studies relying on the timing of the confidence measurement is not consistent with all the existing studies. It has been argued that asking observers to indicate their choice and their confidence at the same time interferes with the confidence report [14]. For example, asking participants to report confidence and choice at the same time might be sufficient to induce a report strategy that is no longer based on posterior probabilities, but on heuristics [36]. Additionally, measuring confidence after the choice may allow observers to collect additional evidence after the choice or even change their minds [3,41,42,46,47]. In favour of the timing-based explanation, those studies to report a decrease of confidence with discriminability assessed first the choice and confidence only after the choice [14, 41]. The studies to report the opposite pattern more often recorded confidence simultaneously with the response [42, 43]. Nevertheless, at least in the masked orientation discrimination task, the timing of the responses does not provide a satisfying explanation, because an increase of confidence in incorrect choices with discriminability was consistently observed irrespective of whether confidence was assessed at the same time as the choice or afterwards [32]. Future experiments appear necessary to test if the timing of the confidence measurement influences patterns of confidence in the other experimental paradigms.

Is there other empirical support for the hypothesis that confidence is not only based on sensory evidence about the identity of the stimulus, but also on evidence about discriminability? There is evidence that the brain represents estimates of discriminability: A recent neuro-imaging study showed that neural areas in posterior parietal cortex and ventral striatum track sensory reliability independently of the choice [4]. To our knowledge, only one study so far included evidence about discriminability into a formal modelling analysis. In a masked orientation discrimination task, confidence was best explained by a combination of evidence about the identity of the stimulus as well as the general visibility of the stimulus, although the study did not test whether evidence about the identity of the stimulus and visibility were combined in a Bayesian fashion [32]. In contrast, when the double increase pattern was observed in random dot kinematograms, the increase of confidence in errors with discriminability was explained by an influence of decision times of confidence [42, 43]. However, at least in the masked orientation discrimination task, decision times cannot not account for the increase of confidence in errors with discriminability because decision time in incorrect trials was uncorrelated with discriminability [32].

Although more experiments are clearly necessary to investigate the relationship between confidence and decision time, the hypothesis regarding *e*_*d*_ gains some plausibility due to converging evidence that human confidence is informed by many cues. One mechanism may rely on the variability of *e*_*I*_: In a random dot motion discrimination task, confidence depended on the consistency of the random dot motion, although discrimination performance was equated [48]. Additionally, when observers discriminated the average colour of an array of coloured shapes, confidence was not only determined by the distance of the average colour to the category boundary, but was also affected by the variability of colour across the array [49]. A second mechanism may rely on the elapsed time during decision making: In in a global motion discrimination task, the time required to make a decision was varied while the sensory evidence about the motion direction was equated, showing that decision time directly informed confidence [42]. Given that human metacognition appears to make use of such a variety of cues, it seems plausible to us that sensory evidence about discriminability may be involved as well.

## Conclusion

To summarize, the present paper argues that the folded X-pattern can be misleading as a signature of confidence. On theoretical grounds, it can be expected that in many psychophysical tasks, confidence in incorrect choices increases, not decreases with discriminability. On empirical grounds, it must be acknowledged that the folded X-pattern can only be observed for some tasks, while it does hold true for other tasks. Overall, it is not legitimate to identify neural correlates of confidence by assuming a specific signature of confidence a priori. When statistical properties are used to track correlates of confidence, it appears essential to empirically assess the pattern of confidence in each single task using behavioural markers of confidence.

## Material and Methods

All simulations were conducted using the free software R [50]. Each simulated experiment consisted of 4×10^6^ trials.

### Standard model

For the standard model, three sets of simulations were performed. Each simulation started with sampling the stimulus d for each single trial of the simulated experiment. We assumed that the identity of the stimulus was −1 and 1 for 2×10^6^ trials each. Then, we sampled discriminability |d|. For the first set of simulations, we simulated 9 experiments, where the discriminability |d| was sampled from a different distribution for each of the nine experiments:

- discrete uniform distribution with the possible values 0, and 1
- discrete uniform distribution with the possible values 0, 1, and 2
- discrete uniform distribution with the possible values 0, 1, 2, 3, and 4
- continuous uniform distribution with min = 0 and max = 1
- continuous uniform distribution with min = 0 and max = 2
- continuous uniform distribution with min = 0 and max = 4
- gamma distribution with a shape α = 1 and rate β = 2
- gamma distribution with a shape α = 1.5 and rate β = 1.5
- gamma distribution with a shape α = 2 and rate β = 1.

The parameters of the gamma distribution were chosen so that the mean and variance of the distribution matched the discrete uniform distributions.

The second set of simulations with the standard model involved four simulated experiments.|d| was always sampled from a discrete uniform distribution, but we varied the set from which |d| was sampled:

- Possible values were 0, 2, and 4
- Possible values were 0, 1, 2, 3, and 4
- Possible values were 0, ½, 1, …, 4
- Possible values were 0, ¼, ½, …, 4

For the third set of simulations with the standard model, |d| was again always sampled from a discrete uniform distribution. In each of the 4 simulated experiments, there were only two possible values of |d|, one of which was always 0. The other possible value of |d| were 1, 2, 3, and 10, respectively.

Then, for each single trial of the stimulated experiments, the sensory evidence eI was sampled from Gaussian distributions with M = d and σ_*I*_ = 1. The choice ϑ was −1 if e_I_ < 0 and 1 otherwise. The accuracy of the choice was defined as correct when I and ϑ were the same. For each single trial, the posterior probability of a correct choice given the percept and the choice p(A=1| ϑ, e_I_) was calculated using the formulae S1 Appendix.

### General model

For the simulation based on the general model, we simulated 36 experiments, one for each combination of 9 possible distributions from which the discriminability d was drawn, and 4 possible levels of noise σ_d_ with respect to the sensory evidence e_d_ about the discriminability. In each experiment, we first sampled the identity of the stimulus I ∈ {−1,1} for each single trial of the experiment. It was assumed that both identities of the stimulus I ={−1,1} occurred 2×10^6^ times. Then, the discriminability d was drawn for each trial of the experiment. We used the same distributions as in the first set of simulations for the standard model. Then, for each single trial of the experiment, the evidence about the identity of the stimulus e_I_ was sampled from Gaussian distributions with M = d × I and σ = 1. When e_I_ was greater than zero, observers were assumed to make the choice ϑ = 1, and ϑ = −1 otherwise. When the choice matched the identity of the stimulus, the choice was considered correct. The evidence about discriminability e_d_ was sampled from Gaussian distributions with M = d and the standard deviation of σ_d_. σ_d_ varied across experiments with the possible values 1/10, 1/3, 1, and 10.

Finally, confidence c was calculated for each single trial as the posterior probability of a correct choice given the sensory evidence for identity, sensory evidence for discriminability, and choice p(A=1| ϑ, e_d_, e_I_) was calculated using the formulae S2 Appendix.

### Accuracy of confidence

The information entropy of choice accuracy conditioned on confidence H(A|c) can be calculated as

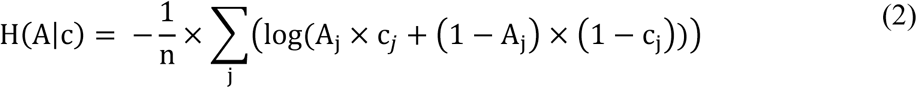

where n is the number of simulated trials, A_j_ is the accuracy in trial j, and c_*j*_is the confidence in trial j.

## Acknowledgements

We are grateful to Christina Linner and Florian Sprang for helpful comments on a previous version of this paper.

## Data availability statement

Reproduceable computer code and all results reported in the present study are available online at the open science framework website (https://osf.io/pfsqz/).

## Supporting information

### S1 Appendix. Derivation of the formula of objective confidence according to the standard model

According to the standard model, it is assumed that an observer selects a choice ϑ ∈ {−1,1} about the identity I ∈ {−1,1} of stimulus d. The identity equals the sign of d, which is sampled in each trial either from a discrete set or from a continuous distribution. The accuracy A ∈ {0,1} of the choice is defined to be 1 if ϑ = I and 0 otherwise. Observers cannot perceive d directly; instead, observers make their choices based on the sensory evidence eI, a noisy estimate of d.

Given the model specification, the posterior probability of being correct given the sensory evidence p(A = 1|e_I_) can be calculated as the posterior probability that identity is the same as the selected choice option, given the sensory evidence. In the following, we consider the case that the observer decides that the identity is 1; formulae for the decision that the identity is −1 can be derived just in the same way.

According to Bayes’ rule, p(I = 1|e_I_) can be calculated as:

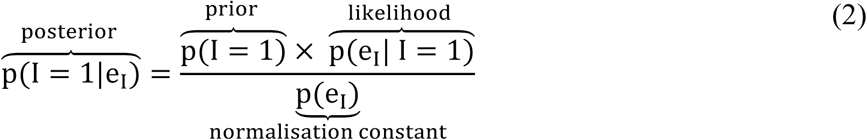

Based on the law of total probability, the normalization constant p(e_I_) can be expressed as:

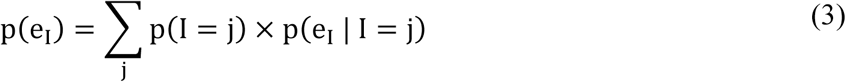

For the purpose of the present analysis, we assumed that the two choice options are equally likely, i.e. the prior probabilities p(I = −1) and p(I = 1) are both 0.5. Therefore, (2) and (3) can be combined and simplified to:

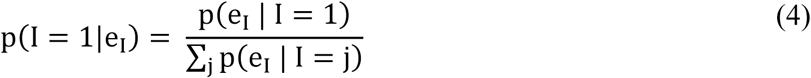

If d is sampled from a finite set of n elements, the denominator of the fraction in (4) can be expressed as a sum of the likelihood of the sensory evidence conditioned on d over all possible values of d weighed by the probability of the specific d. For the numerator, the sum takes into account only positive values of d:

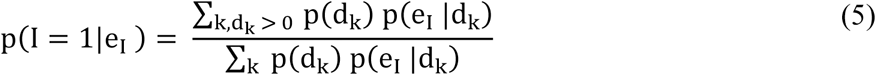

If d is sampled from a continuous distribution instead, numerator and denominator of the fraction in (4) can be expressed as integrals over d:

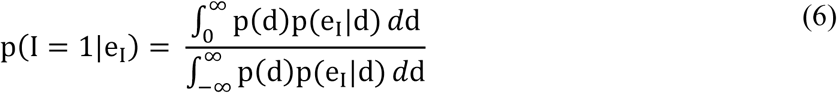

In (6), *d* denotes the differential, while d denotes the stimulus.

### S2 Appendix. Derivation of the formula of objective confidence according to the general model

According to the general model, the observer is presented with a series of stimuli characterised by two features, the identity I ∈ {−1,1} and discriminability d. It is assumed that observers in each trial make a choice ϑ ∈ {−1,1} about the identity I ∈ {−1,1} of the stimulus. The accuracy A ∈ {0,1} of the choice is 1 if ϑ = I and 0 otherwise. The decision is based on sensory evidence, which involves evidence about the stimulus strength e_d_, which depends only on d, and evidence about the identity e_I_, which depends on d and on I.

Given the model specification, the posterior probability of being correct given the sensory evidence p(A = 1|(e_d_, e_I_)) can be calculated as the posterior probability that the identity I of the stimulus is the same as the selected choice option, given the sensory evidence. In the following, we consider the case that the observer decides that I is 1; the formula for the choice that I is −1 can be derived analogously.

According to Bayes’ rule, p(I = 1|(e_D_, e_I_)) can be calculated as:

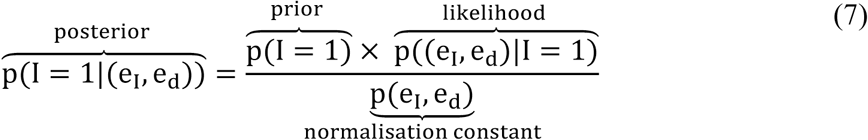

As we assumed again that the two choice options are equally likely and thus the prior probabilities of both identities are the same, formulae (7) can be simplified analogously to formula (6):

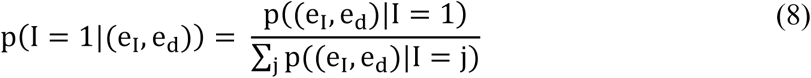

If d is sampled from a discrete set of elements, numerator and denominator in (8) can be expressed as a sum of likelihoods of the sensory evidence conditioned on d over the different values of d. The sum is weighed by the probability of each specific d:

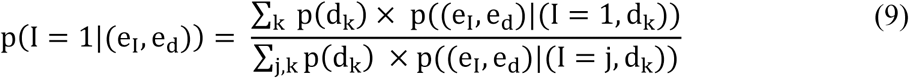

Given the case that d is sampled from a continuous distribution, p(I = 1|(e_I_, e_d_)) can be obtained by integration:

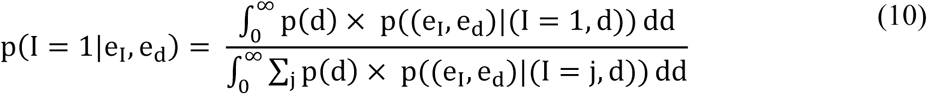

Again, *d* denotes the differential, while d denotes the discriminability of the stimulus. As we assume that e_I_ and e_d_ are stochastically independent when the stimulus strength d is controlled, the likelihood p((e_I_, e_d_)|(I, d)) can be calculated as:

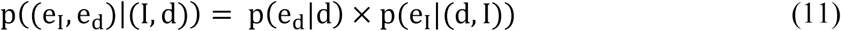

We insert formula (11) into (9) to calculate p(I = 1|e_i_, e_D_) in the discrete case.

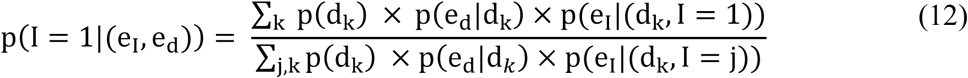

Finally, we insert formula (11) into (10) to calculate p(I = 1|(e_I_, e_d_)) in the continuous case.

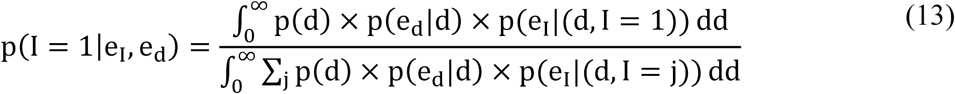

